# Large-scale GWAS reveals genetic architecture of brain white matter microstructure and genetic overlap with cognitive and mental health traits (n=17,706)

**DOI:** 10.1101/288555

**Authors:** Bingxin Zhao, Jingwen Zhang, Joseph G. Ibrahim, Tianyou Luo, Rebecca C. Santelli, Yun Li, Tengfei Li, Yue Shan, Ziliang Zhu, Fan Zhou, Huiling Liao, Thomas E. Nichols, Hongtu Zhu

## Abstract

**Background:** Individual variations of white matter (WM) tracts are known to be associated with various cognitive and neuropsychiatric traits. Diffusion tensor imaging (DTI) and genome-wide single-nucleotide polymorphism (SNP) data from 17,706 UK Biobank participants offer opportunity to identify novel genetic variants of WM tracts and explore the genetic overlap with other brain-related complex traits.

**Method:** We analyzed the genetic architecture of 110 tract-based DTI parameters, carried out genome-wide association studies (GWAS) and performed post-GWAS analyses, including association lookups, gene-based association analysis, functional gene mapping, and genetic correlation estimation.

**Results:** DTI parameters are substantially heritable for all WM tracts (mean heritability 48.7%). We observed a highly polygenic architecture of genetic influence across the genome (p-value=1.67*10^−05^) as well as the enrichment of genetic effects for active SNPs annotated by central nervous system cells (p-value=8.95*10^−12^). GWAS identified 213 independent significant SNPs associated with 90 DTI parameters (696 SNP-level and 205 locus-level associations; p-value<4.5*10^−10^, adjusted for testing multiple phenotypes). Gene-based association study prioritized 112 significant genes, most of which are novel. More importantly, association lookups found that many of the novel SNPs and genes of DTI parameters have previously been implicated with cognitive and mental health traits.

**Conclusions:** The present study identifies many new genetic variants at SNP, locus and gene levels for integrity of brain WM tracts and provides the overview of pleiotropy with cognitive and mental health traits.

## Introduction

Complex brain functions rely on dynamic interactions between distributed brain areas operating in large-scale networks. Consequently, the integrity of white matter connections between brain areas is critical to proper function. Microstructural differences in white matter (WM) tracts are phenotypically associated with information processing speed and intelligence (1–5) as well as neurodegenerative/neuropsychiatric traits, such as Alzheimer’s disease (6), Parkinson’s disease (7), schizophrenia (SCZ) (8), and attention-deficit/hyperactivity disorder (ADHD) (9). A better understanding of genetic factors influencing integrity of WM tracts could have important implication for understanding the etiology of these diseases as well as individual variation in intelligence. To reveal the underlying genetic contributions to brain structural development and disease/disorder processes, imaging genetics studies of WM microstructure has been an active research area over the past fifteen years. The structural changes of WM tracts are typically measured and quantified in diffusion tensor imaging (DTI) (10). Brain diffusivity can be influenced by many aspects of its micro- or macro-structures (11). To reconstruct the WM pathways and tissue microstructure, DTI models the diffusion properties of WM using random movement of water. Specifically, DTI quantifies diffusion magnetic resonance imaging (dMRI) in a tensor model and analyzes diffusions in all directions. A typical DTI diagonalizes the tensor and calculates three pairs of eigenvalues/eigenvectors that respectively represent one primary and two secondary diffusion directions. Within each voxel, several DTI parameters can be derived: fractional anisotropy (FA), mean diffusivity (MD), axial diffusivity (AD), radial diffusivity (RD), and mode of anisotropy (MO). As a summary measure of WM integrity (12, 13), higher FA indicates stronger directionality in this voxel. MD quantifies the magnitude of absolute directionality, AD is the eigenvalue of the principal direction, RD is the average of the eigenvalues of the two secondary directions, and MO is the third moment of a tensor. Positive MO reflects narrow tubular water diffusion, whereas a negative value denotes planar water diffusion (14). There are several approaches to analyze DTI data across the whole brain, including manual region-of-interest (ROI) analysis, automated ROI analysis, voxel-based analysis, such as tract-based spatial statistics (TBSS) (15), as well as tractography and graph theory analysis; see Tamnes, Roalf (16) for a survey.

In family-based studies, the magnitude of genetic influences (i.e., heritability) in various DTI parameters of WM tracts, including FA, MD, AD, and RD, has been examined across a wide age range, from neonates (17, 18), young children (19), older children (20, 21), adolescents (22), and young adults (23) to middle aged (24) and older adults (25). Participants in these studies are typically monozygotic and dizygotic twins or family members. Table 1 of Vuoksimaa, Panizzon (24) lists 14 studies that illustrated that a substantial proportion of variance in DTI parameters (FA, MD, AD, and RD) was explained by additive genetic effects. However, the genetic architecture of DTI parameters remains largely unknown due to the limitation of family-based studies, for which the heritability estimation has relied on contrasting the phenotypic similarity between monozygotic and dizygotic twins. Genetic architecture denotes the characteristics of genetic variations that contribute to the broad-sense heritability of a phenotype (26). Based on the number of genetic variants contributing to phenotypic variance, genetic architecture can be described as monogenic (one variant), oligogenic (few variants), polygenic (many variants), or omnigenic, which hypothesizes that almost all genetic variants have small but non-zero genetic contributions (27, 28). Uncovering the genetic architecture and discovering the associated genetic variants are essential steps to delineate the functional mechanisms and understand the genetic overlap between white matter structures and neuropsychiatric traits.

Recent developments have enabled heritability estimation and genetic variants discovery with using the common single-nucleotide polymorphisms (SNPs) data collected in general populations. Instead of using the expected genetic correlation based on pedigree information, SNP heritability is estimated by adding up the genetic effects across a large number of common SNPs (minor allele frequency [MAF]>0.05 or 0.01) (29, 30). The architecture of genetic influences can be assessed by SNP annotation and partition (31, 32). Genome-wide association studies (GWAS) and post-GWAS analysis can further identify causal genetic variants at SNP, locus and gene levels (33, 34), and assess the genetic overlap of complex traits in different domains (35, 36). With these methods, the availability of genomic and imaging data from recent large population-based United Kingdom (UK) Biobank resource (37) offers the opportunity to uncover the genetic basis of brain WM tracts in one large-scale, relatively homogeneous population. The UK Biobank has captured data from over 500,000 original participants of middle or elderly ages (age range 40-69), and is currently in the process of following up with 100,000 of these participants to perform brain MRI screening (38).

Here we generated 110 tract-based DTI parameters using the British ancestry UKB sample including 17,706 participants. For each of the 110 phenotypes, we estimated the SNP-heritability, assessed the distribution of genetic effects by SNP annotation and partition, and carried out GWAS to identify the associated genetic variants at SNP and locus levels. We further discovered gene-level associations via MAGMA (39), and explored the functional consequences of the significant SNPs by functional mapping and annotation analysis (FUMA, (34)). To detect genetic overlap and pleiotropy in WM tracts and other complex traits, we performed association lookups at SNP and gene levels on the NHGRI-EBI GWAS catalog (40) and estimated genetic correlations via LD score regression (LDSC, (36)). The GWAS summary statistics have been made publicly available at https://med.sites.unc.edu/bigs2/data/gwas-summary-statistics/.

## Materials and Methods

### Participants and image preprocessing

We used data from 17,706 UKB individuals of British ancestry (self-reported ethnic background, Data-Field 21000). The DTI data (38) and covariates were downloaded from the UKB data resource. We generated 110 DTI parameters: FA, AD, MD, MO and RD of 21 WM tracts, and their average values across these tracts. The ID and full names of the 21 WM tracts are listed in **Supplementary Table 1.** A full description of the DTI registration and quality controls is documented in supplementary information and an overview is given in **Supplementary Figure 1**. We removed values greater than five times the median absolute deviation from the median in each continuous variable. All individuals were aged between 40 and 80 years and the proportion of females was 52.9%.

### Genotyping and quality control

We downloaded the imputed SNP data from UKB data resource (41). We further performed the following SNP data quality controls using PLINK (42): excluding subjects with more than 10% missing genotypes, only including SNPs with MAF > 0.01, genotyping rate > 90%, and passing Hardy-Weinberg test (p-value>1*10^−7^). We also removed SNPs with imputation INFO score less than 0.8.

### SNP-heritability analysis and genome-wide association analysis

For each of the 110 DTI parameters, we estimated the proportion of variation explained by all autosomal SNPs with using linear mixed-effects model (LMM) via GCTA (29). We considered the fixed effects of age (at imaging), age-squared, gender, age-gender interaction, age-squared-gender interaction, as well as the top 40 genetic principal components (GPCs, Data-Field 22009). We also estimated the proportion of variation explained by SNPs in each chromosome. In addition, we partitioned the SNPs according to cell-type-specific annotations (32) to perform enrichment analysis. SNPs were grouped according to their functional activeness in various cell groups and specifically in the central nervous system (CNS) cell group: CNS_active, CNS_inactive, and Always_inactive, see supplementary information for detailed definitions. We performed GWAS for each DTI parameter separately with PLINK (42). The same set of covariates were adjusted in all heritability and GWAS analyses.

### Genomic risk loci characterization and comparison with previous findings

We characterized genomic risk loci by using FUMA (34) online platform (v1.3.4). FUMA first identified independent significant SNPs, which were defined as significant SNPs that were independent of each other (R^2^<0.6). FUMA then constructed LD block for independent significant SNPs by tagging all SNPs that had a MAF ≥ 0.0005 and were in LD (R^2^≥0.6) with at least one of the independent significant SNPs. If LD blocks of independent significant SNPs were closed (<250 kb based on the closest boundary SNPs of LD blocks), they were merged to a single genomic locus. More details of FUMA analysis can be found in (34). Independent significant SNPs and all SNPs in LD with them were subsequently searched on NHGRI-EBI GWAS catalog (40) (v2019-01-31) to look for reported associations (p-value<9*10^−6^) with any traits.

### Gene-based association analysis and functional annotation

We carried out gene-based association analysis for 18,796 protein-coding candidate genes via MAGMA (39) (v1.07). Gene-based p-values were calculated by summarizing the GWAS results of corresponding SNPs, which were mapped to genes according to their psychical positions. Significant genes were searched on NHGRI-EBI GWAS catalog (40) (v2019-01-31) to look for their previously reported associations with any traits. We focused on brain-related complex traits and characterized them into six groups: cognitive (e.g., general cognitive ability, cognitive performance, math ability, and intelligence), education (e.g., years of education and college completion), reaction time, neuroticism, neurodegenerative diseases (e.g., Alzheimer’s disease, Parkinson’s disease and corticobasal degeneration), and neuropsychiatric disorders (e.g., major depressive disorder [MDD], SCZ, bipolar disorder [BD], ADHD, alcohol use disorder, and autism spectrum disorder).

We also performed functional gene annotation and mapping via FUMA. SNPs were annotated with their biological functionality and then were linked to genes by a combination of positional, expression quantitative trait loci (eQTL) association, and 3D chromatin interaction mappings. Specifically, independent significant SNPs and all SNPs in LD with them were annotated for gene functional consequences by ANNOVAR (43). The annotated SNPs were mapped to 35,808 candidate genes based on physical position on the genome (tissue/cell types for 15-core chromatin state: brain), eQTL associations (tissue types: GTEx v7 brain (44), BRAINEAC (45), and CommonMind Consortium (46)) and chromatin interaction mapping (built-in chromatin interaction data: dorsolateral prefrontal cortex, hippocampus (47); annotate enhancer/promoter regions: E053-E082 brain (48)). We used default values for all other parameters in FUMA.

### Genetic correlation estimation with LDSC

We used LDSC (v1.0.0, https://github.com/bulik/ldsc) to estimate the pairwise genetic correlation between DTI parameters and other traits by their GWAS summary statistics. In LDSC, we used the pre-calculated LD scores provided by LDSC (https://data.broadinstitute.org/alkesgroup/LDSCORE/), which were computed using 1000 Genomes European data. We used HapMap3 SNPs and removed all SNPs in chromosome 6 in the MHC region.

## Results

### SNP heritability estimation

**Figures 1-2** and **Supplementary Figures 2-7** display the SNP heritability of DTI parameters estimated by all common autosomal SNPs. The associated standard errors, raw and Bonferroni-adjusted p-values from the one-sided likelihood ratio tests are given in **Supplementary Table 2**. All SNP heritability estimates were significantly larger than zero (Bonferroni-adjusted p-value<0.004). Genetic factors accounted for a moderate or large portion of the variance of DTI parameters in all WM tracts (mean heritability 0.487, standard errors are around 0.041). For example, genetic effects explained more than 60% of the total variance of FA in the posterior limb of the internal capsule (PLIC), anterior corona radiata (ACR), superior longitudinal fasciculus (SLF), and cingulum cingulate gyrus (CGC). SNP heritability of FA decreased to 37% and 27%, respectively, in the fornix (FX) and corticospinal tract (CST). According to the functions of the WM tracts (Connectopedia Knowledge Database, http://www.fmritools.com/kdb/white-matter/), we clustered them into four communities: complex fibers (C1), associative fibers (C2), commissural fibers (C3) and projection fibers (C4). We found that the set of WM tracts in C1 and C3 (mean=0.512) tended to have higher SNP heritability than those in C2 and C4 (mean=0.440, p-value=2.16*10^−04^).

**Figure 1.**
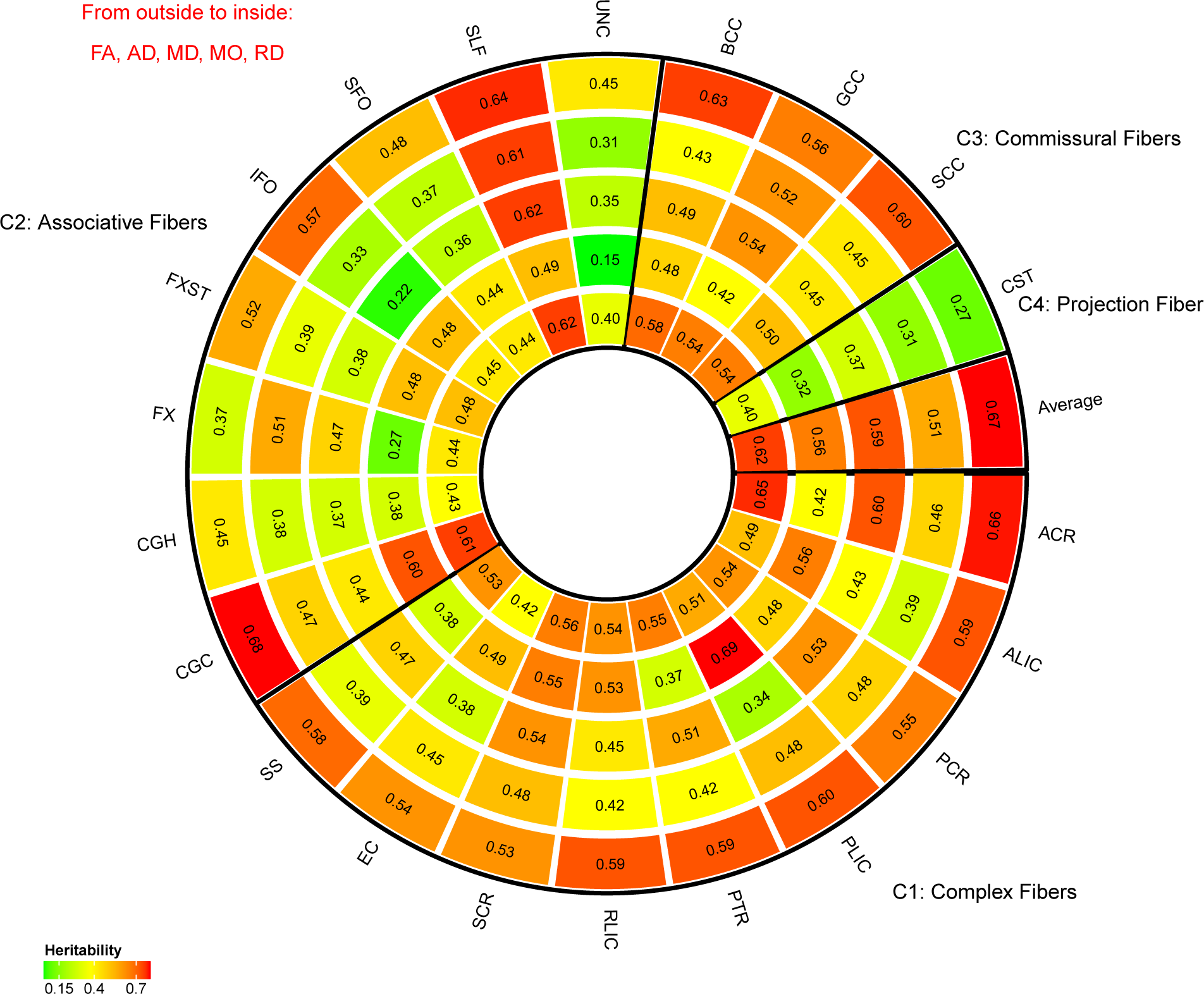
SNP heritability estimates grouped by white matter tract functions.

**Figure 2.**
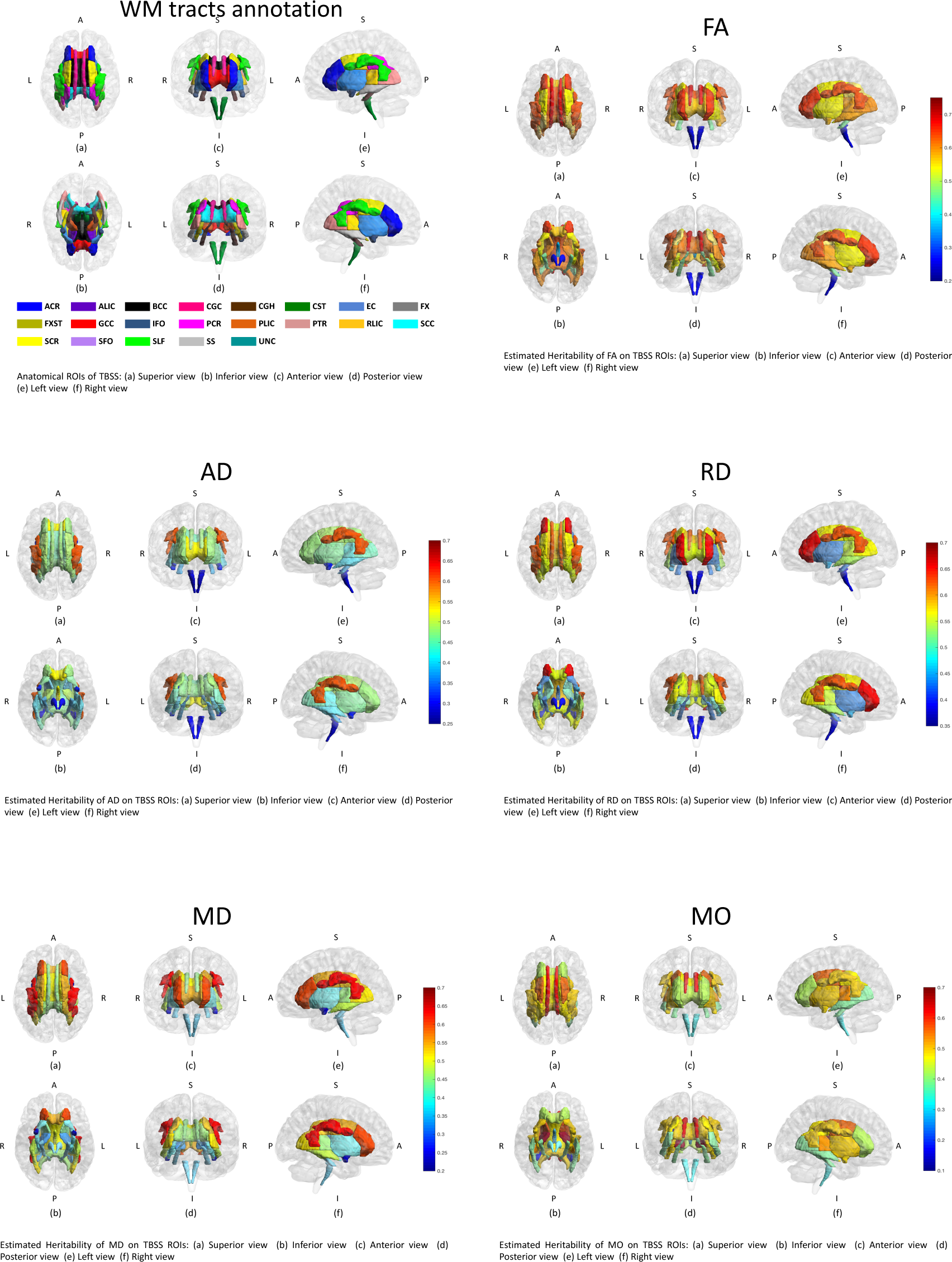
Distribution of SNP heritability estimates of the 21 white matter tracts in brain.

### Partitioning and annotating genetic variation

To examine the distribution of SNP heritability across the genome, we partitioned SNP data into 22 chromosomes and estimated the SNP heritability by each chromosome **(Supplementary Table 3)**. We found that the mean heritability across all 110 DTI parameters explained by each chromosome was linearly associated with the length of the chromosome **(Figure 3(a)**, R^2^=61.2%, p-value=1.67*10^−05^). This finding reveals a highly polygenic or omnigenic genetic architecture (28) of WM tracts. The large number of SNPs that contribute to the variation in DTI parameters are widely spread across the whole genome. To further illustrate this architecture, we ordered and clustered the 22 chromosomes into three groups by their lengths: long, medium, and short. The long group had 4 chromosomes (CHRs 2, 1, 6, 3), which together accounted for 33% of the length of the whole genome; the medium group had 6 chromosomes (CHRs 4, 5, 7, 8, 10, 11), which accounted for another 33% of the length of the whole genome; and the short group consisted of the remaining 12 chromosomes. **Figure 3(b)** shows the SNP heritability estimates grouped by chromosomal length. It is clear that longer chromosomes tended to have higher SNP heritability estimates than medium (p-value=3.82*10^−13^) or shorter (p-value<2.20*10^−16^) ones for DTI parameters.

**Figure 3.**
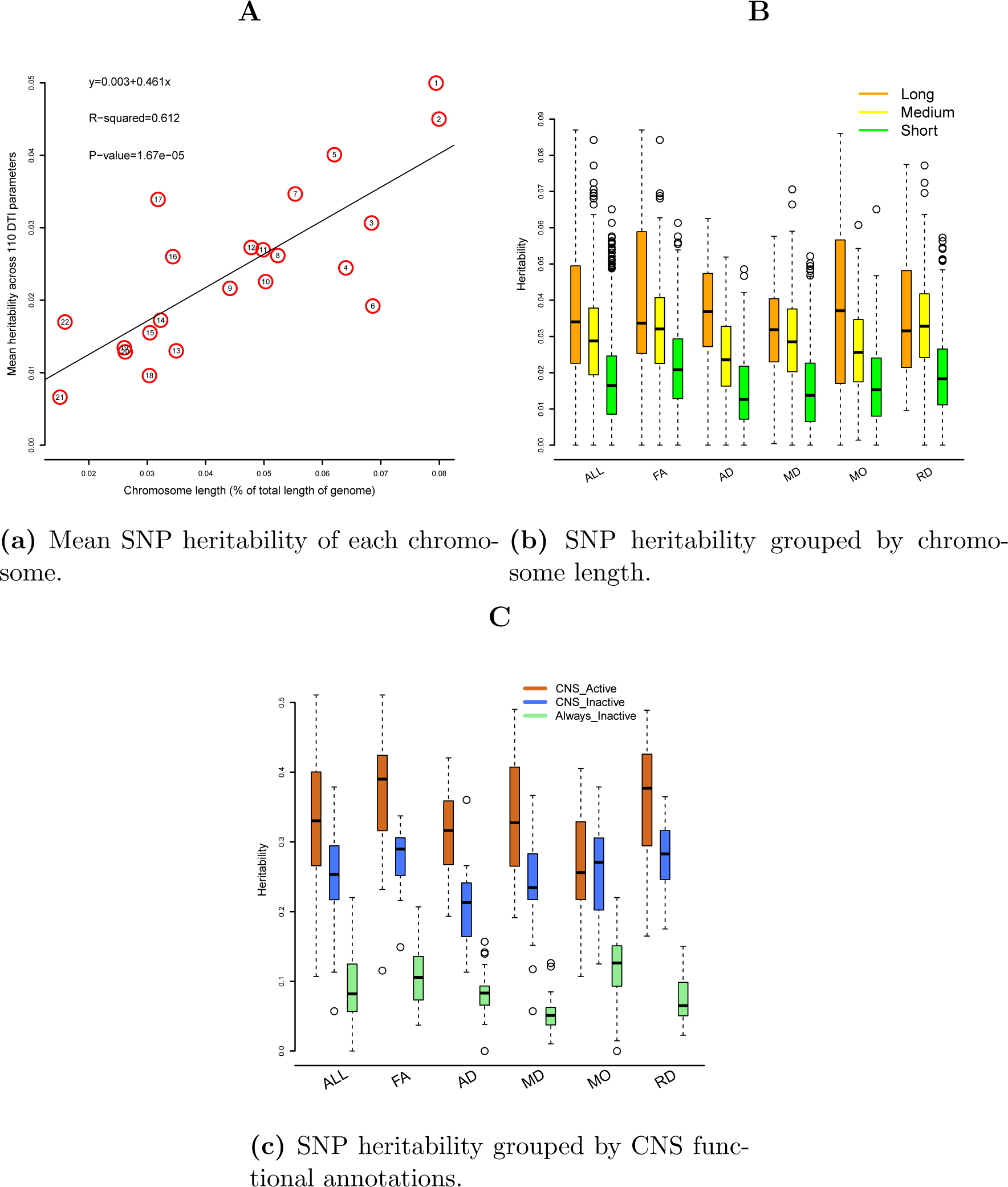
Heritability estimated by SNPs in each chromosome or in functionally annotated SNP categories.

To compare the contribution of SNPs with different activity level, we partitioned the genetic variation according to CNS-cell-specific annotations: CNS_active, CNS_inactive, and Always_inactive **(Supplementary Table 4)**. Heritability estimated by SNPs residing in chromatin regions inactive across all cell groups (Always_inactive) was clearly much smaller than the heritability estimated by SNPs residing in chromatin regions active in CNS cell (CNS_active, p-value<2.20*10^−16^). The heritability estimated by CNS_inactive SNPs (inactive in CNS cell but active in other cells) was also significantly smaller than that of CNS_active SNPs (p-value=8.95*10^−12^) **(Figure 3(c))**. This pattern remained consistent across all the five types of DTI parameters, though larger variance was observed for the MO parameters.

### GWAS results of 110 DTI parameters

We carried out GWAS for the 110 DTI parameters with using 8,955,960 SNPs after genotyping quality controls. All Manhattan and QQ plots are shown in **Supplementary Figure 8**. 19,530 significant associations were detected at the 4.5*10^−10^ significance level (that is, 5*10^−8^/110, adjusted for testing multiple phenotypes) **(Supplementary Figure 9, Supplementary Table 5)**. RD and MD of anterior limb of internal capsule (ALIC) had more than 3,000 significant associations. Significant SNPs were summarized into 213 independent significant SNPs by FUMA, which had 696 independent significant associations with 90 DTI parameters **(Figure 4, Supplementary Tables 6-7**). RD and FA of splenium of corpus callosum (SCC) had the largest number of independent significant SNPs. 502 of the 696 independent significant associations located in chromosome 5 **(Supplementary Table 8, Supplementary Figure 10)**. The 696 independent significant SNP-level associations can be further characterized as 205 locus-level associations **(Supplementary Table 9)**. FA and RD of SCC, FA and AD of FX, and RD of ALIC had at least five genetic risk loci **(Supplementary Table 10)**. Each chromosome had at least one genetic risk locus except for chromosomes 13,20 and 21, and chromosome 5 had the largest number of risk loci **(Supplementary Tables 11)**. GWAS results at 5*10^−9^ and 5*10^−8^ significance levels are also reported in above tables and figures.

**Figure 4.**
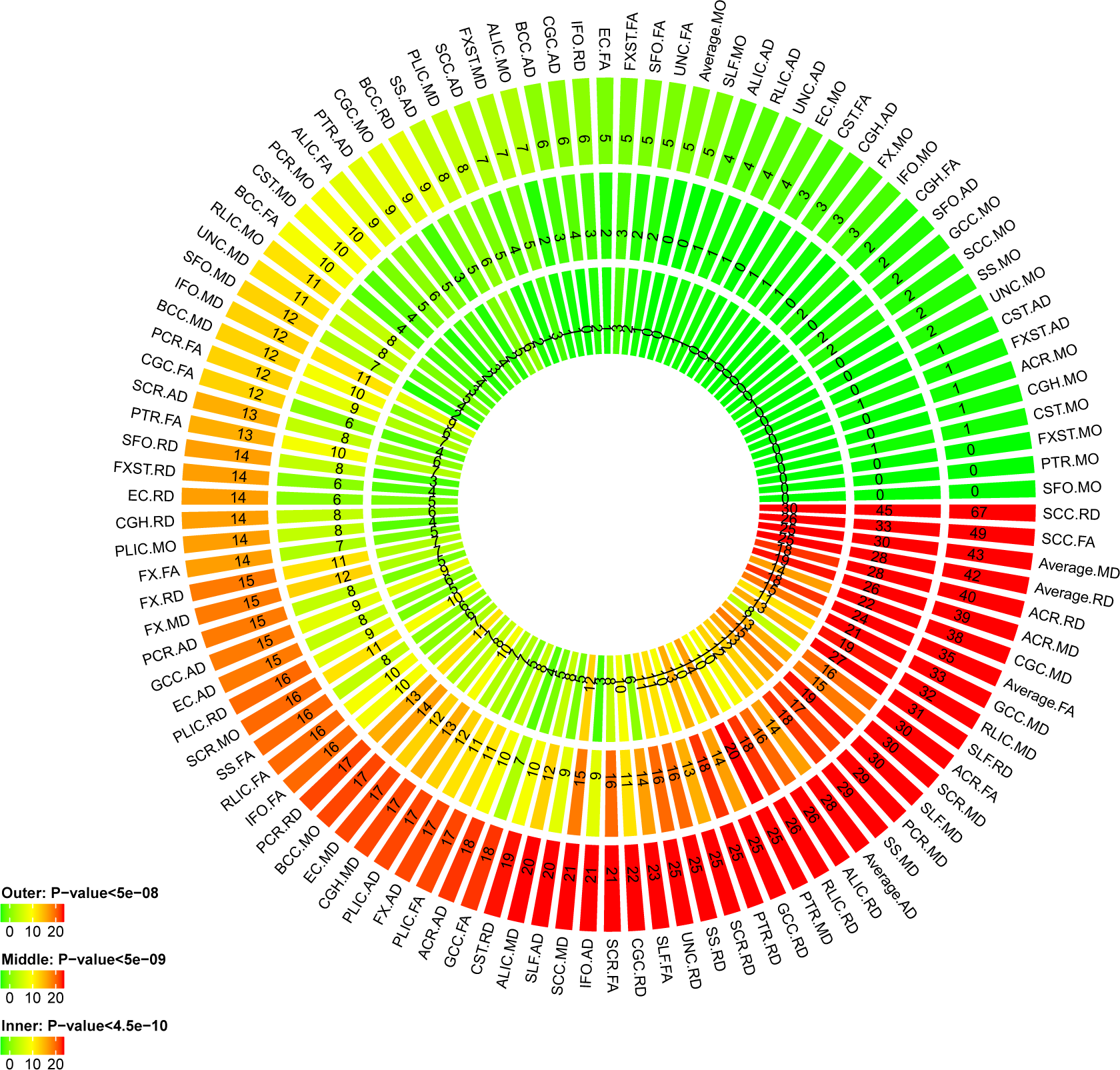
Number of independent significant SNPs discovered for each DTI parameter at different GWAS significance levels. Outer layer: p-value <5*10^−8^; middle layer: p-value <5*10^−9^; and inner layer: p-value <4.5*10^−10^.

### Concordance with previous GWAS results

Association lookups on the NHGRI-EBI GWAS catalog (40) found that 122 of the 213 independent significant SNPs (associated with 83 DTI parameters) were reported to be associated with any traits **(Supplementary Table 12)**. Our study replicated many SNPs reported in previous GWAS of WM hyperintensity measures and other brain structural measures **(Supplementary Table 13)**, most of which were recently detected in Rutten-Jacobs, Tozer (49) (n=8,448). In addition, we tagged 15 different SNPs associated with neuropsychiatric disorders, 40 with cognitive traits, 12 with education, 47 with neuroticism, 17 with neurodegenerative diseases, and 2 with reaction time. We also compared our results with the those reported in Elliott, Sharp (50) (n=8,428) and found that 212 of the 368 significant associations (**Supplementary Table 6** of (50)) were replicated in the present study. We note that the both (49) and (50) analyzed a subset of the sample presented in this study.

### Gene-based association analysis and functional mapping

Gene-based association analysis identified 508 significant gene-level associations (p-value<2*10^−8^, adjusted for testing multiple phenotypes) between 112 genes and 96 DTI parameters **(Supplementary Table 14).** Our results replicated genes discovered in Rutten-Jacobs, Tozer (49) and Elliott, Sharp (50), including *VCAN, C16orf95, NBEAL1, SH3PXD2A, CACNB2, SRA1, GNA12, CPED1*, and *EPHA3,* but most of the identified genes were not previously linked to DTI parameters. Association lookups found that 51 of the 112 significant genes were implicated with cognitive, education, reaction, neuroticism, neuropsychiatric and neurodegenerative traits in previous studies, such as *CRHR1* (51-54), *MAPT* (55-58), *KANSL1* (59-61), and *MSRA* (62–64) **(Supplementary Table 15, Figure 5)**. We also annotated the SNPs by functional consequences on gene functions **(Supplementary Figure 11)** and performed functional gene mapping. Gene mapping discovered 292 genes **(Supplementary Table 16)**, 218 of which were not detected in the gene-based association analysis.

**Figure 5.**
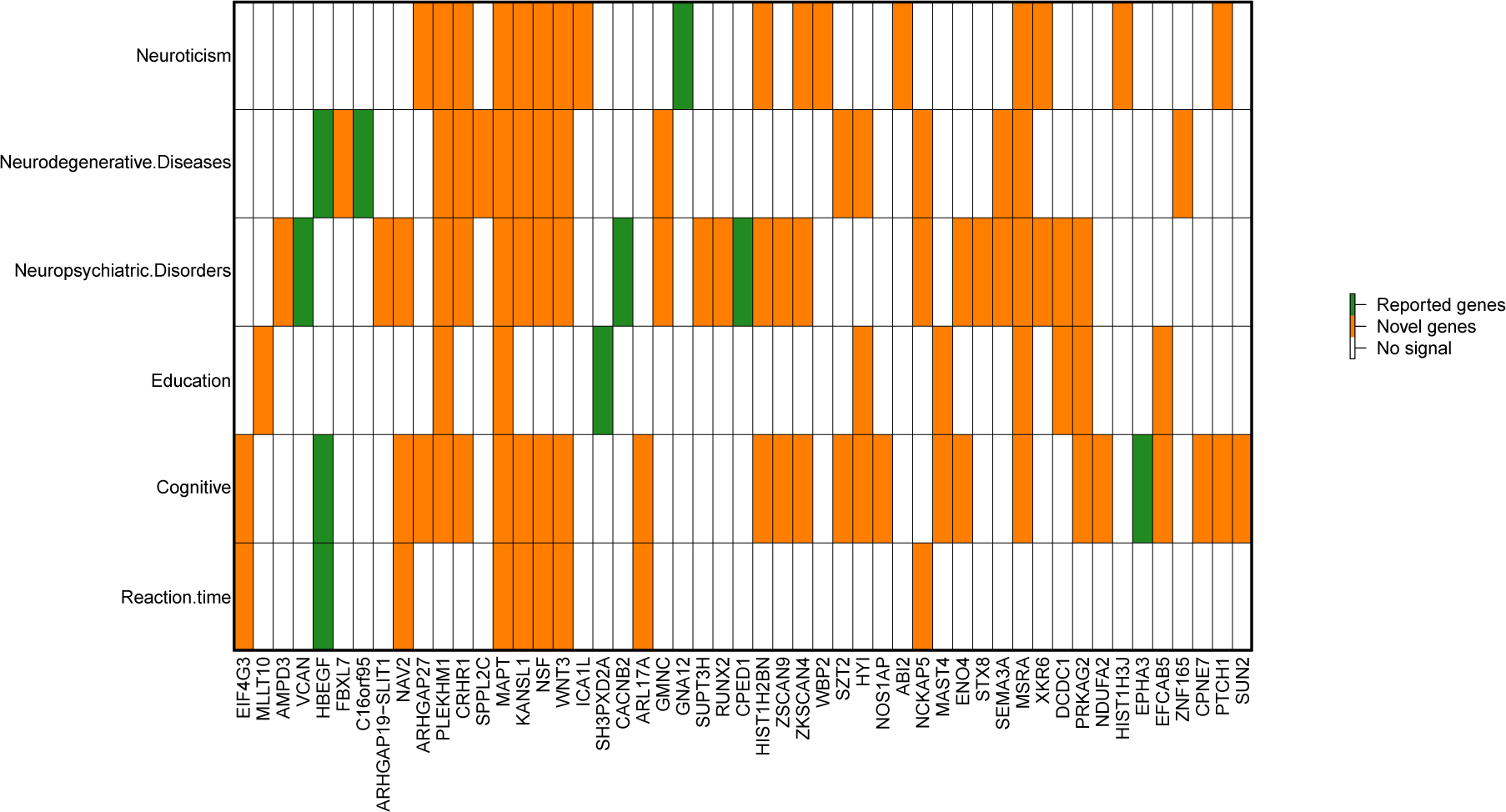
Genes identified in gene-based association analysis of DTI parameters that have been implicated with traits of neuroticism, neurodegenerative diseases, neuropsychiatric disorders, education, cognitive, and reaction time in previous GWAS.

### Genetic correction with other traits

We estimated the pairwise genetic correlation between 110 DTI parameters and 14 other complex traits **(Supplementary Table 17)**. We focused on traits showing evidence of pleiotropy in association lookups. 43 pairs of phenotypes had significant genetic correlation after adjusting for multiple testing (1,540 tests) by the Benjamini-Hochberg (B-H) procedure (65) at 0.05 level **(Supplementary Table 18, Supplementary Figure 12).** Reaction time had significant negative correlations with FA parameters (mean=-0.181), and had widespread positive correlations with AD, MO, MD and RD (mean=0.165) **(Figure 6)**. Education, cognitive, intelligence, and numerical reasoning also had positive genetic correlations with AD, FA, and MO. On the other hand, depression, MDD and drink frequency showed negative genetic correlations with FA. Other pairs were insignificant after multiple testing adjustment.

**Figure 6.**
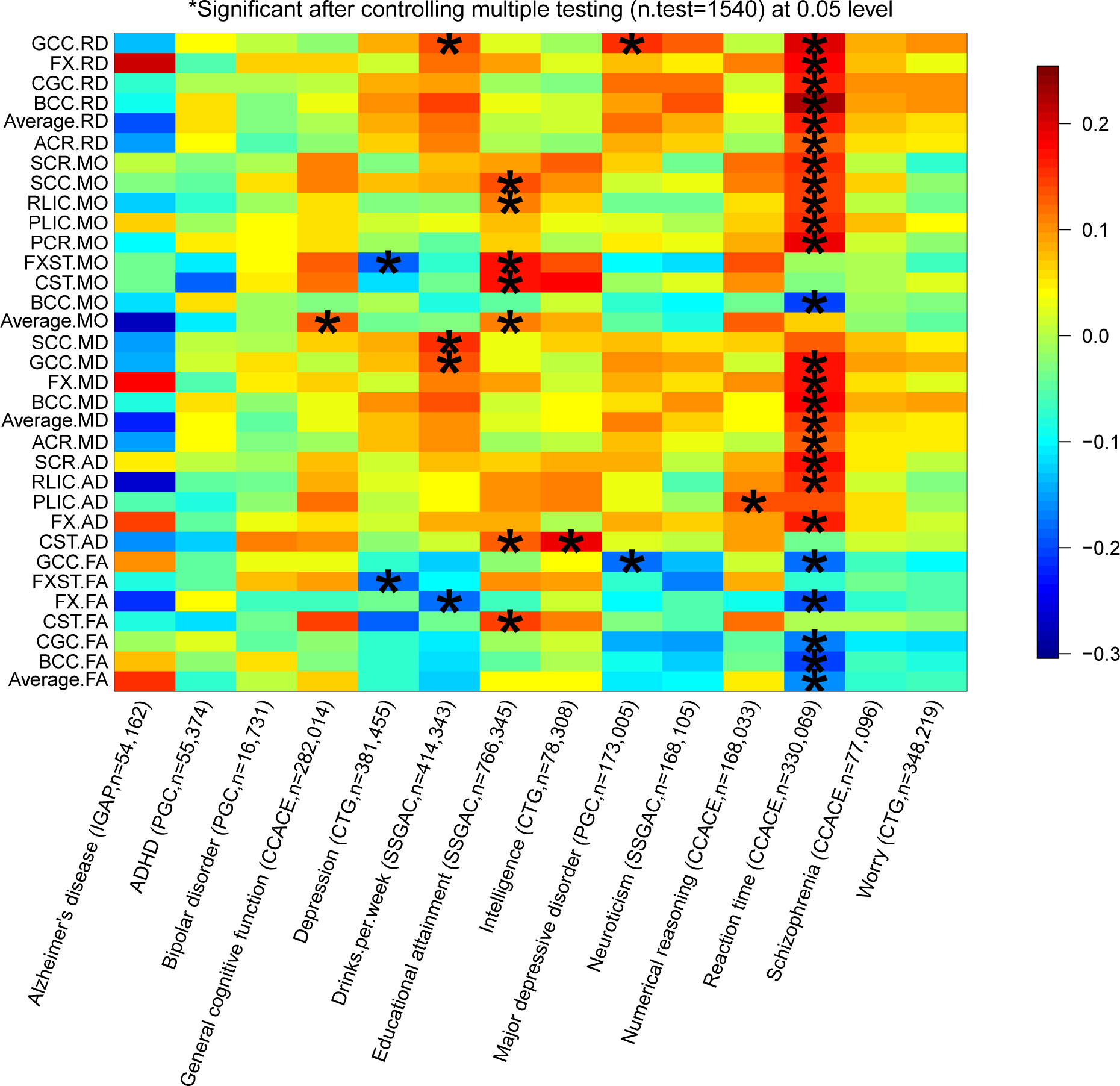
Selected pairwise genetic correlations between DTI parameters and other traits.

## Discussion

Heritability and GWAS analyses can provide guidance for downstream analyses to model the functional mechanisms and pathways involved in the phenotype of interest or its pleiotropy traits. A large number of family-based neuroimaging studies have documented that WM tracts are essentially heritable across the lifespan. Two recent GWAS (49, 50) have made attempts to explore the genetic risk variants of DTI parameters, however, they were less powered due to the limited sample size (n<9,000). With the DTI and common autosomal SNP data from 17,706 UKB participants, the present study made novel contributions to 1) understand the genetic landscape of WM tract; 2) identify novel genetic risk variants at SNP, locus, and gene levels; and 3) provide the overview of statistical pleiotropy (35, 66) with other brain-related complex traits.

Our SNP heritability estimates are close to the ones reported in previous family-based studies (e.g., Table 1 of (24)), and are also within a similar range as those reported in (50), where the mean heritability is around 0.450. These results suggest that studies of DTI phenotypes using common SNPs may be more informative than studies focused on rare variants. Our results partitioning the genetic variation in chromosomes or SNP functional sets shed light on the distribution of genetic signals across the genome and different functional consequences. These findings suggest a highly polygenic genetic architecture of DTI parameters and also provide evidence for stronger genetic signals from SNPs in active chromatin regions, especially for those active in the CNS cell type. For such highly polygenic traits, large sample size is essential for GWAS to discover the widespread genetic signals (67). Compared to previous GWAS (49, 50), our study with larger sample size identifies many newly associated genetic variants for DTI parameters. More importantly, these novel findings uncover the widespread pleiotropy between DTI parameters and cognitive and metal health traits. As the UKB releases more imaging data, it can be expected that better powered genetic studies on heritable WM tracts will continue facilitating gene exploration and helping understand the causal relationships of brain-related complex traits.

Our analyses reflect several methodological limitations of the current approaches on population-based imaging genetic studies. First, similar to previous studies (23), CST and FX were reported to have low SNP heritability, which may be due to the fact that such small, tubular tracts cannot be well registered and reliably resolved with current techniques (68). Second, heritability estimated by SNP data reflects narrow-sense heritability, which only considers the additive genetic effects of common variants. The genetic architecture may change as we broadly consider all genetic contributions (such as rare variants, non-additive effects and gene-gene interactions) in future studies. However, it is notable that with common SNPs in the UK Biobank, we have gained heritability estimates comparable to those reported in family-based studies. Finally, it is worth mentioning that the UKB data used in this study were sampled from a specific cohort (British ancestry) with a specific age-range. Since genetic ancestries are common confounding effects and aging can play an important role in brain WM structure changes, one should be careful to generalize these findings to general populations or to specific clinical cohorts. With more data from diverse imaging genetics studies, future research will be required to overcome these limitations and advance our biological understanding of the human brain.

## Funding

This research was partially supported by U.S. NIH grants MH086633 and MH116527, and a grant from the Cancer Prevention Research Institute of Texas.

## Acknowledgements

We thank the individuals represented in the UK Biobank dataset for their participation and the research teams for their work in collecting, processing and disseminating these datasets for analysis. This research has been conducted using the UK Biobank resource (application number 22783), subject to a data transfer agreement. We gratefully acknowledge all the studies and databases that made their GWAS summary data available. The authors acknowledge the Texas Advanced Computing Center (TACC, http://www.tacc.utexas.edu) at The University of Texas at Austin for providing HPC and storage resources that have contributed to the research results reported within this paper.

## Conflict of interest

The authors declare no competing financial interests.

