## Supplementary_information for "Large-scale GWAS reveals genetic architecture of brain white matter microstructure and genetic overlap with cognitive and mental health traits (n=17,706)"

#### Supplementary figures

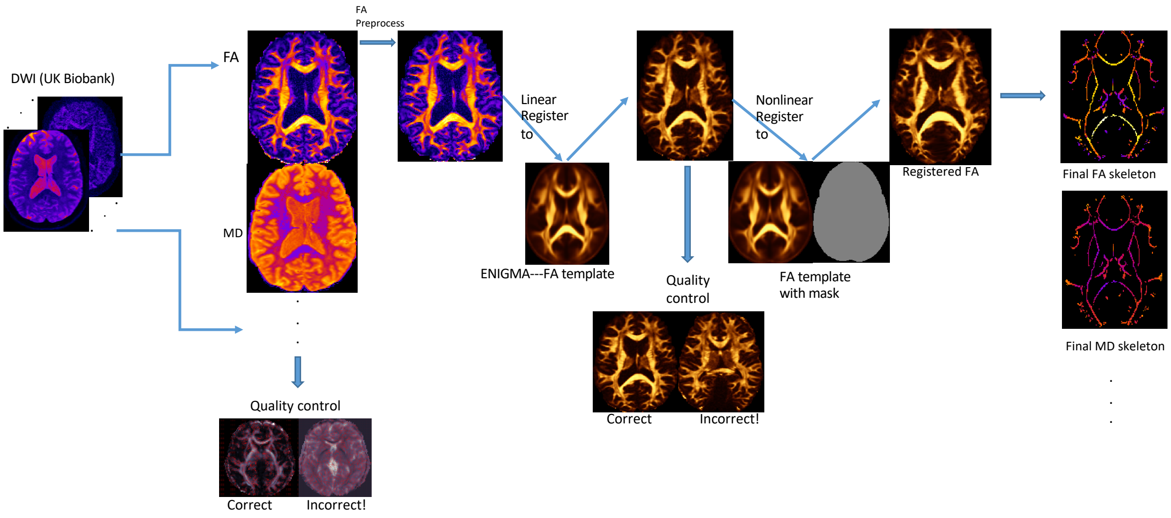


**Supplementary Figure 1:** Overview of the ENIGMA-DTI pipeline for DTI processing.

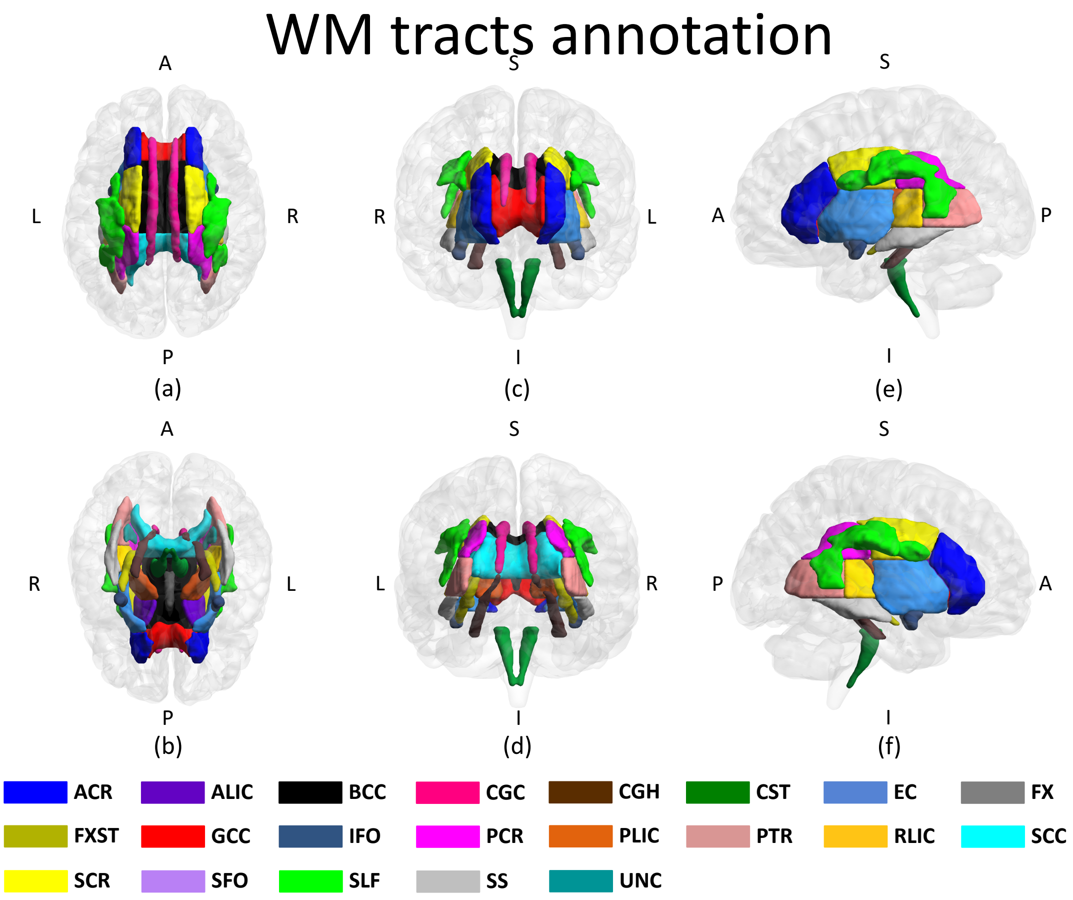


Anatomical ROIs of TBSS: (a) Superior view (b) Inferior view (c) Anterior view (d) Posterior view (e) Left view (f) Right view

**Supplementary Figure 2:** 21 WM tracts annotation.

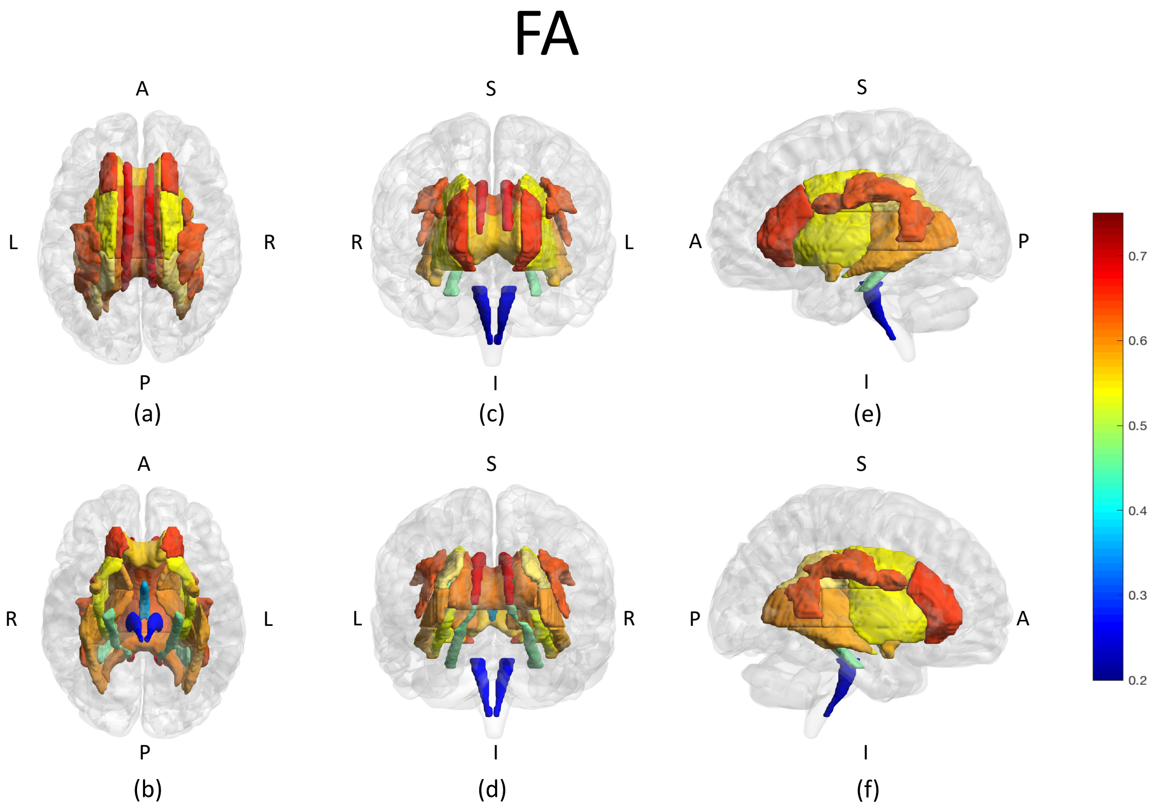


Estimated Heritability of FA on TBSS ROIs: (a) Superior view (b) Inferior view (c) Anterior view (d) Posterior view (e) Left view (f) Right view

**Supplementary Figure 3:** SNP heritability estimates of FA for 21 WM tracts.

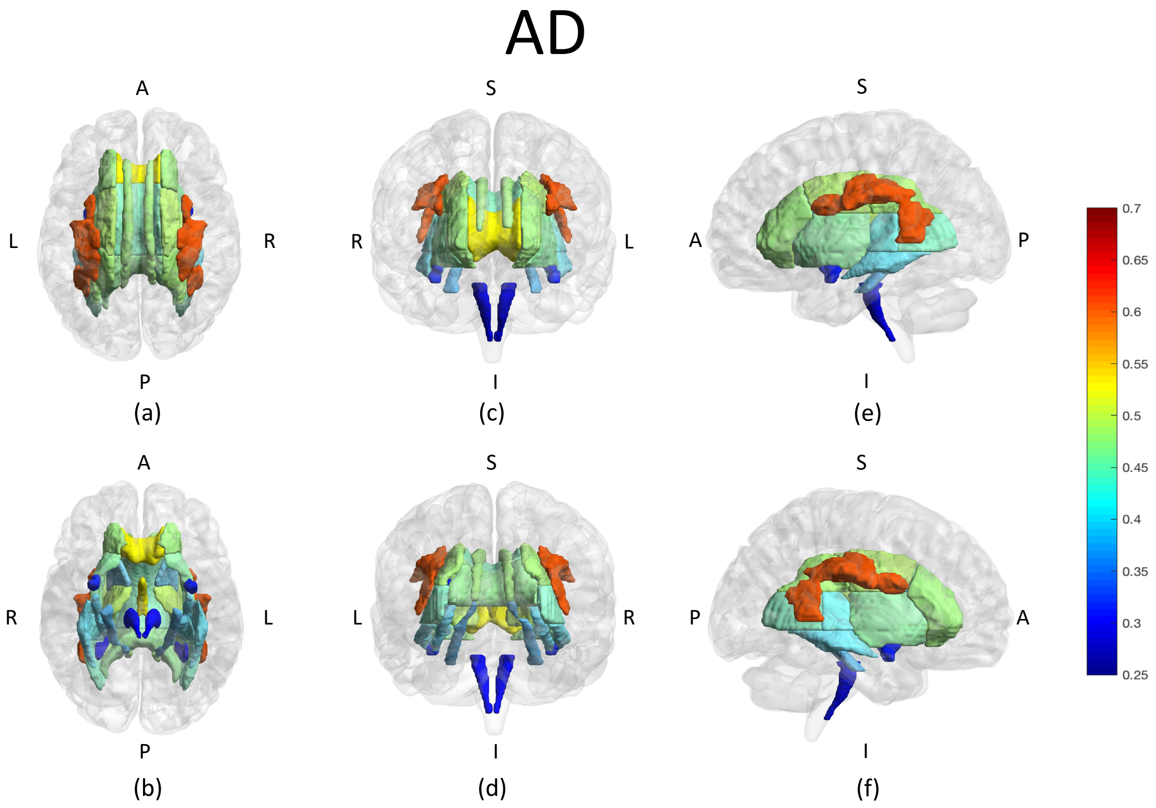


Estimated Heritability of AD on TBSS ROIs: (a) Superior view (b) Inferior view (c) Anterior view (d) Posterior view (e) Left view (f) Right view

**Supplementary Figure 4:** SNP heritability estimates of AD for 21 WM tracts.

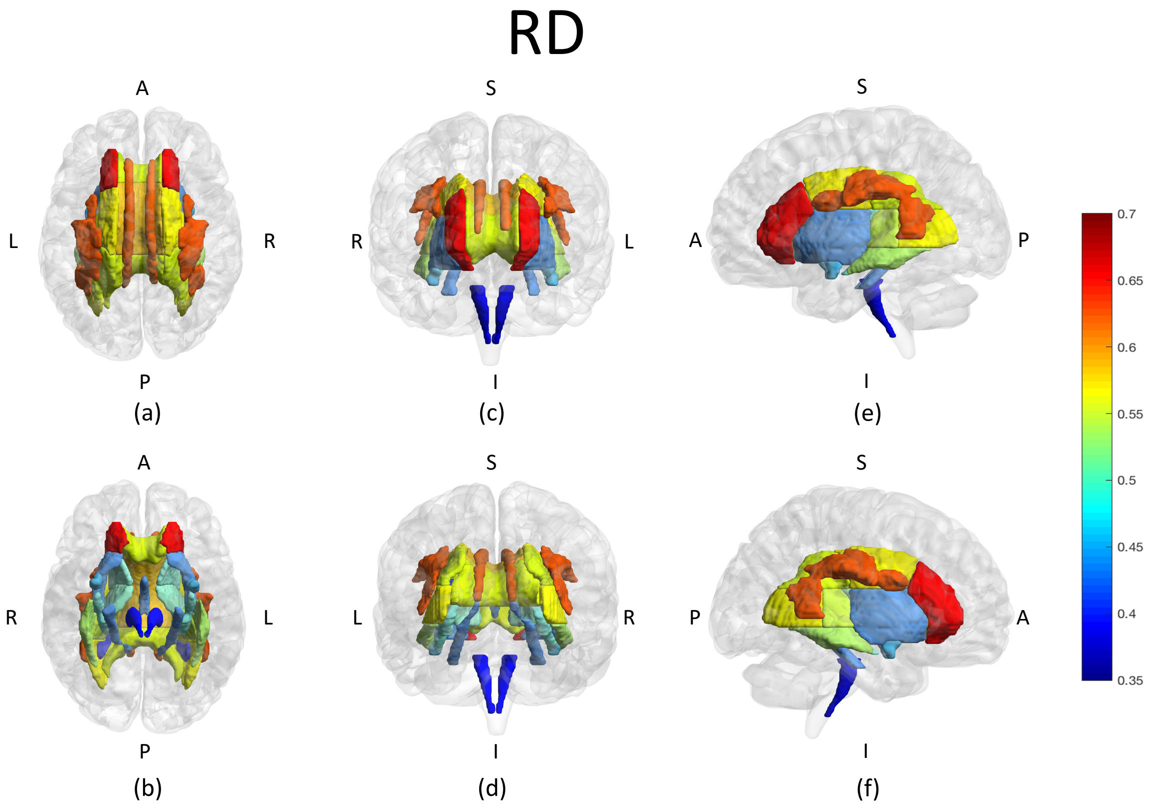


Estimated Heritability of RD on TBSS ROIs: (a) Superior view (b) Inferior view (c) Anterior view (d) Posterior view (e) Left view (f) Right view

**Supplementary Figure 5:** SNP heritability estimates of RD for 21 WM tracts.

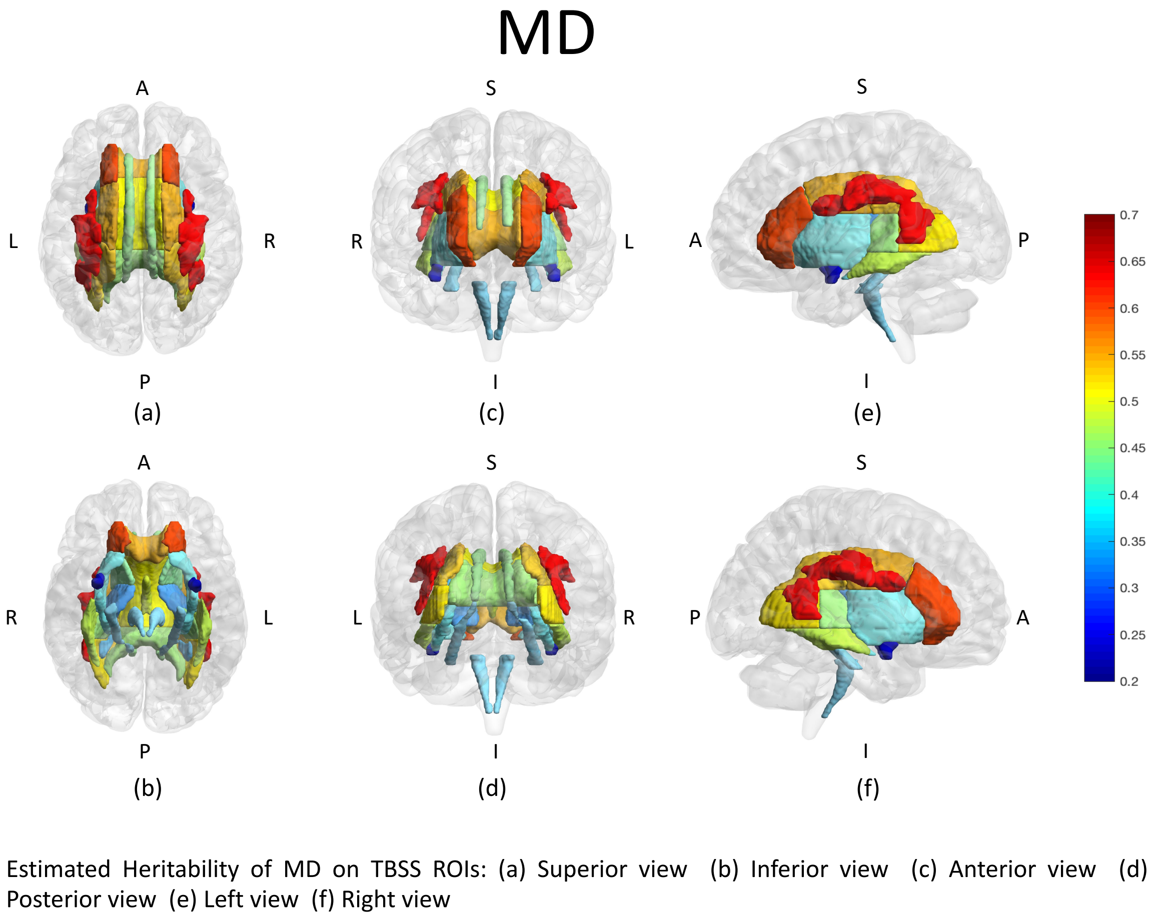


**Supplementary Figure 6:** SNP heritability estimates of MD for 21 WM tracts.

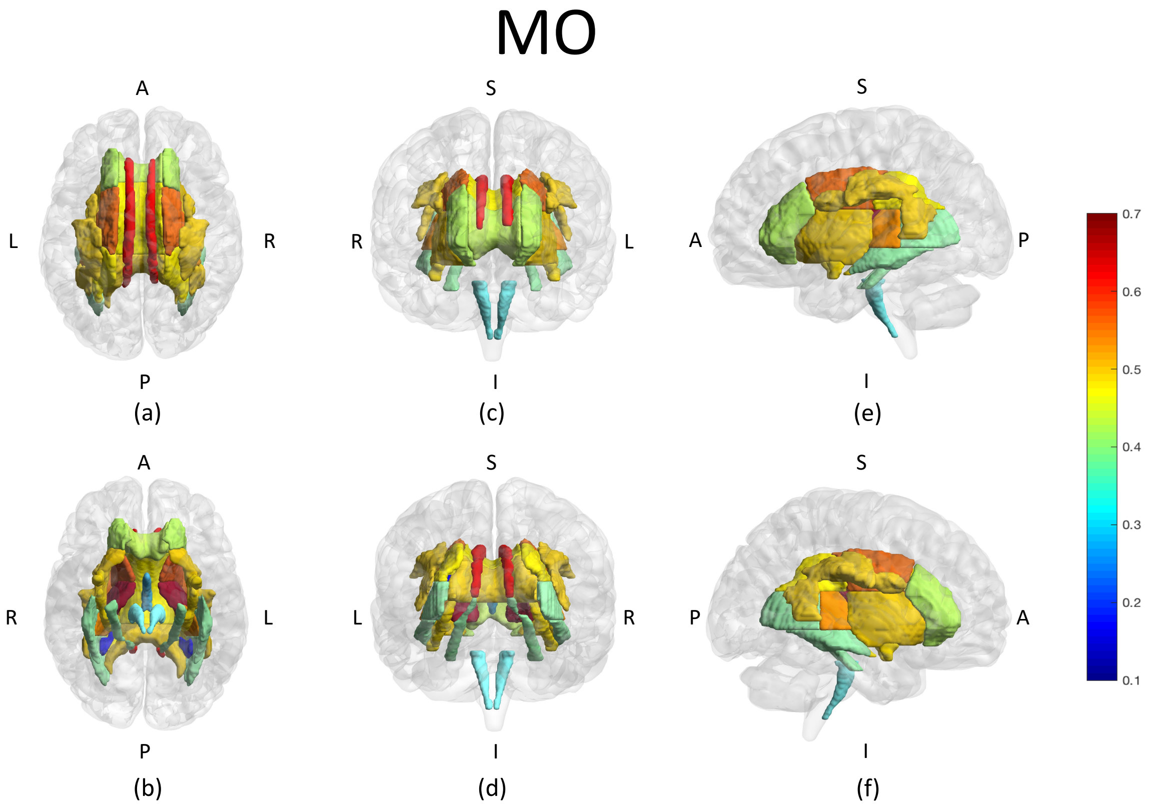


Estimated Heritability of MO on TBSS ROIs: (a) Superior view (b) Inferior view (c) Anterior view (d) Posterior view (e) Left view (f) Right view

**Supplementary Figure 7:** SNP heritability estimates of MO for 21 WM tracts.

**Supplementary Figure 8:** GWAS Manhattan and QQ plots for 110 DTI parameters, available at `https://www.dropbox.com/s/zp89vjcle85uxu3/ukbiobank_dti110_figures.zip?dl=0`

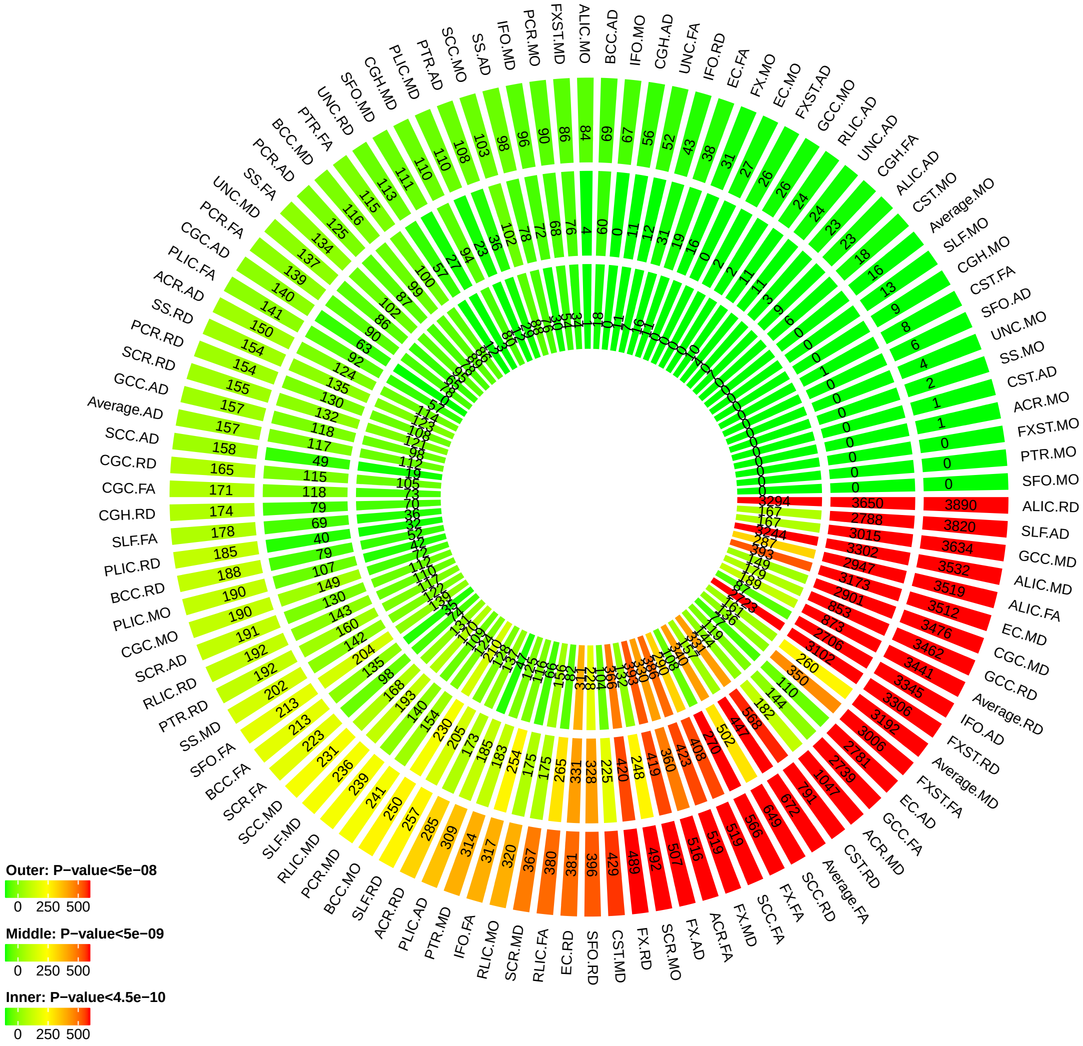


**Supplementary Figure 9:** Number of significant SNP associations discovered at different significance levels. Outer layer: P-value $< 5 \times 10^{-8}$; middle layer: P-value $< 5 \times 10^{-9}$; and inner layer: P-value $< 4.5 \times 10^{-10}$.

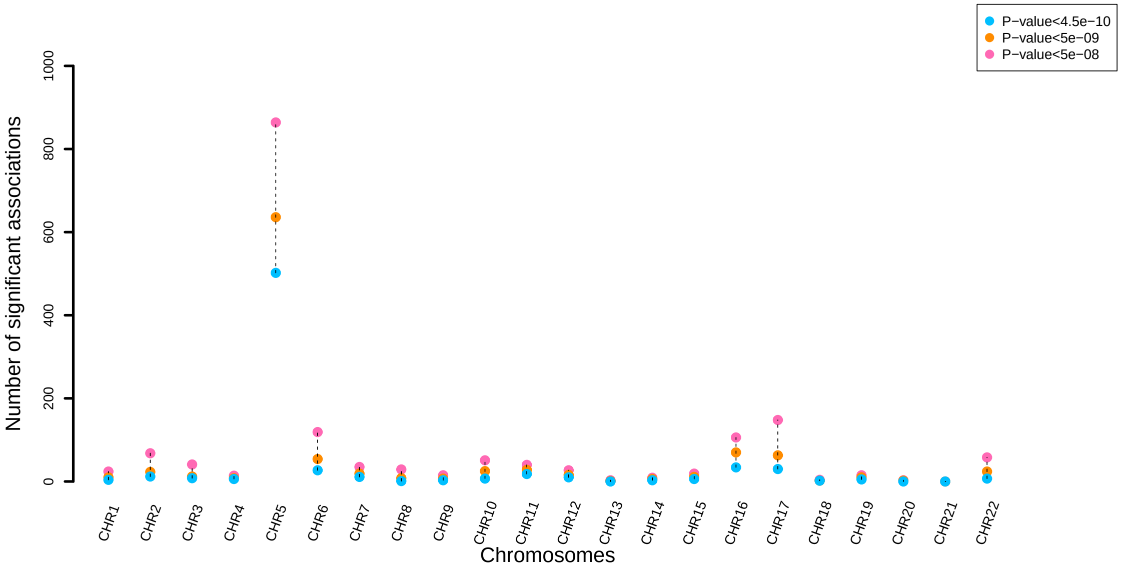


**Supplementary Figure 10:** Number of independent significant SNP associations for each chromosome at different significance levels.

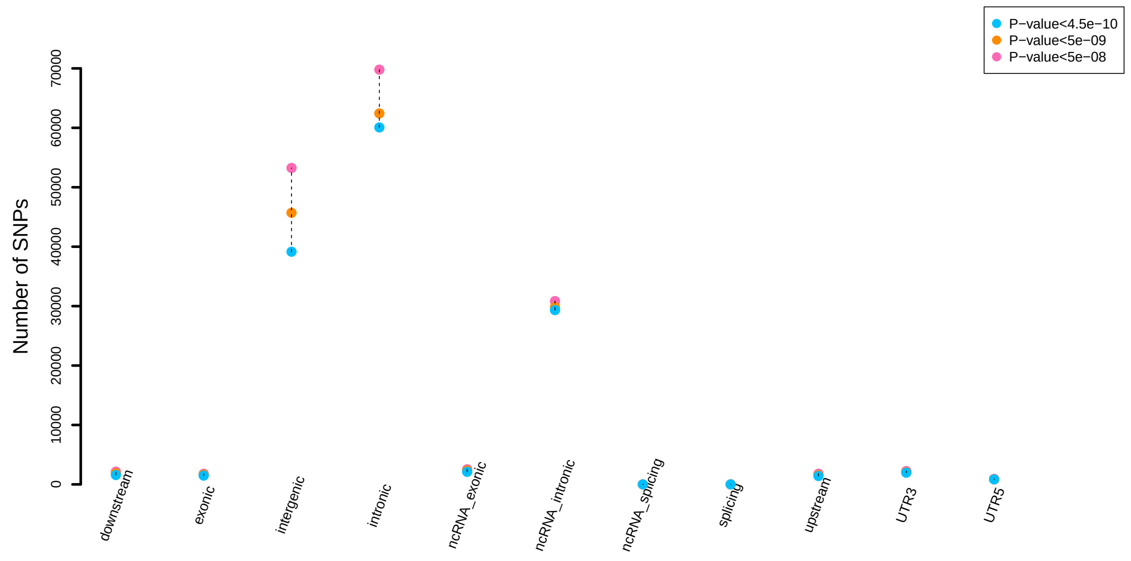


**Supplementary Figure 11:** Functional consequences of indepdenent SNPs (and SNPs in LD with them) indicated by functional annotation assigned by ANNOVAR at different significance levels.

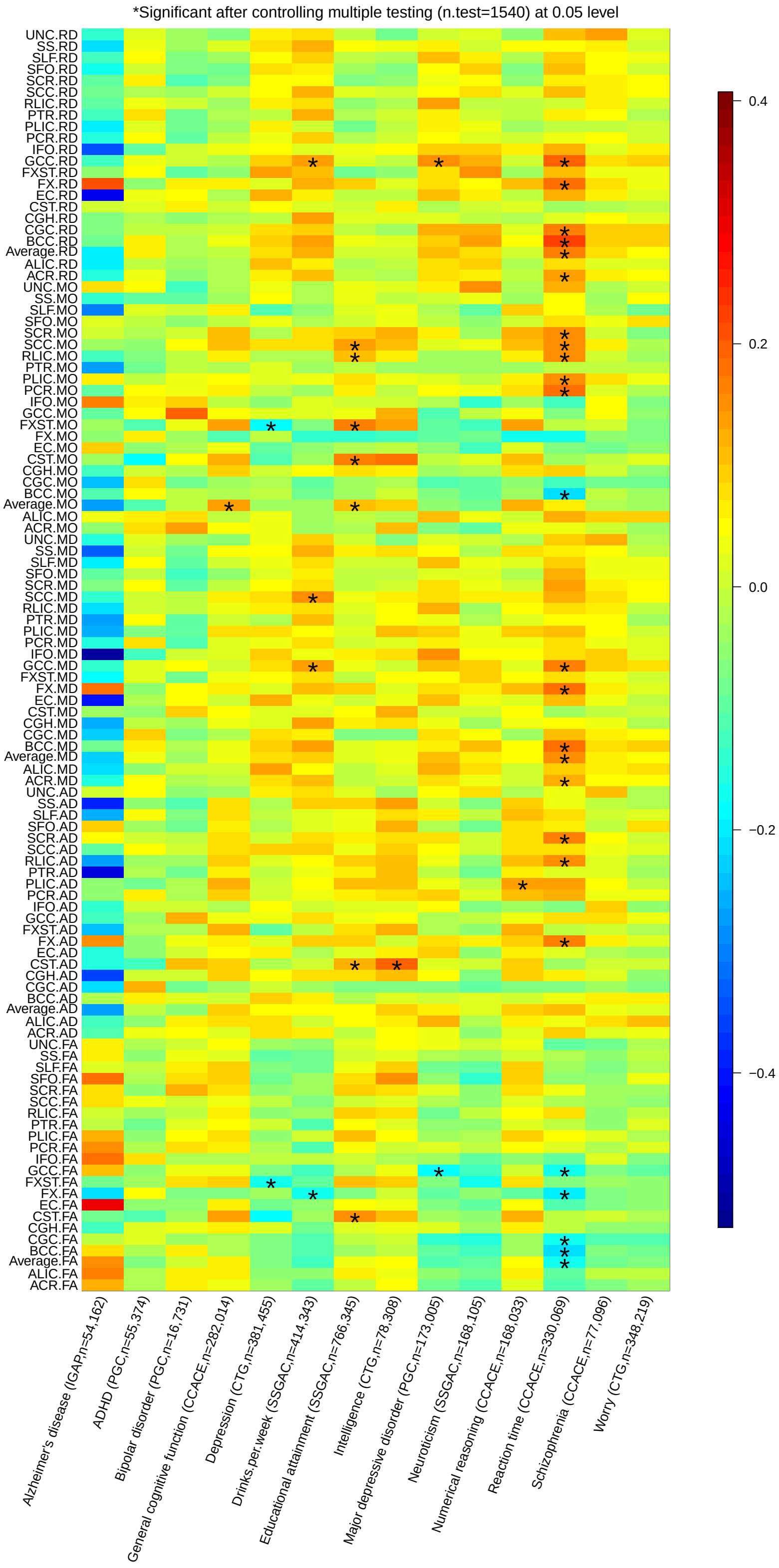


**Supplementary Figure 12:** Pairwise genetic correlation between 110 DTI parameters and other traits. Stars are significant associations after adjusting for multiple testing by the Benjamini-Hochberg procedure at 0.05 level.

### 6 Supplementary tables

**Supplementary Table 1:** List of IDs and full names of the white matter tracts analyzed in this study.

**Supplementary Table 2:** SNP heritability of DTI parameters estimated by all common autosomal SNPs.

**Supplementary Table 3:** SNP heritability of DTI parameters estimated by SNPs in each chromosome.

**Supplementary Table 4:** SNP heritability of DTI parameters estimated by SNPs residing in different chromatin regions.

**Supplementary Table 5:** Number of significant SNP associations at different significance levels.

**Supplementary Table 6:** List of independent significant SNP associations at different significance levels.

**Supplementary Table 7:** Number of independent significant SNP associations at different significance levels.

**Supplementary Table 8:** Number of independent significant SNP associations for each chromosome at different significance levels.

**Supplementary Table 9:** List of significant genetic risk loci at different significance levels.

**Supplementary Table 10:** Number of significant genetic risk loci at different significance levels.

**Supplementary Table 11:** Number of significant genetic risk loci for each chromosome at different significance levels.

**Supplementary Table 12:** Independent significant (P-value $< 4.5 \times 10^{-10}$) SNPs and their correlated SNPs of DTI parameters that have previously been identified at P-value $< 9 \times 10^{-6}$ in GWAS of any traits listed in the NHGRI-EBI GWAS catalog (version 2019-01-31, www.ebi.ac.uk/gwas/).

**Supplementary Table 13:** Independent significant (P-value $< 4.5 \times 10^{-10}$) SNPs and their correlated SNPs of DTI parameters that have previously been identified at P-value $< 9 \times 10^{6}$ in GWAS of any DTI parameters and brain white matter and structural traits listed in the NHGRI-EBI GWAS catalog (version 2019-01-31, www.ebi.ac.uk/gwas/).

**Supplementary Table 14:** List of significant gene-level associations identified by MAGMA (P-value $< 2 \times 10^{-8}$).

**Supplementary Table 15:** Associated traits of MAGMA-identified genes that have previously been reported in the NHGRI-EBI GWAS catalog (version 2019-01-31, www.ebi.ac.uk/gwas/).

**Supplementary Table 16:** List of mapped genes by functional mapping of GWAS results at different significance levels.

**Supplementary Table 17:** Sources of the 14 sets of publicly available GWAS summary statistics used in genetic correlation estimation analysis.

**Supplementary Table 18:** Genetic correlation estimates and p-values between DTI parameters and other traits.

### Supplementary Methods

#### Image processing

DTI data of the UK Biobank were acquired at the 2*2*2 mm spatial resolution with multi-band acceleration factor of three (i.e., three slices are acquired simultaneously) and two b-values (b = 1,000 and 2,000 s/mm2), anterior-to-posterior (AP) is the acquisition phase-encoding direction. For each of these b-values, 50 diffusion-encoding directions were acquired, and there were 100 distinct directions in total. The echo time (TE) was 92 ms and the repetition time (TR) was 3600 ms. The diffusion preparation was a standard (mono-polar) Stejskal-Tanner pulse sequence. The shorter echo time (TE = 92 ms) can enable a higher signal-to-noise ratio (SNR) than a twice-refocused (bipolar) sequence at the expense of stronger eddy current distortions (Miller et al., 2016). We downloaded the UKB derived FA, AD, MD, MO, and RD image maps (Alfaro-Almagro et al., 2018) and performed standard registration and quality controls based on the ENIGMA-DTI pipeline (Jahanshad et al., 2013; Kochunov et al., 2014). We applied consistent procedures for each of the five groups of parameters. Specifically, we first used linear registration to register each of the FA images to the MNI (Montreal Neurological Institute) ICBM152 template at 1*1*1 mm spatial resolution. We then applied nonlinear registration to align the linearly registered FA images to the ENIGMA FA template and masked the registered FA with a template mask. Next, we projected the ENIGMA skeleton onto the registered FA images to skeletonize them and extract ROI-based FA statistics. We excluded subjects whose FA images did not pass the standard imaging quality controls (based on directional information from the primary eigenvector and registration performance, see `http://enigma.ini.usc.edu/protocols/dti-protocols/` for details). Finally, the ROI-based statistics of other four types of DTI parameters (AD, MD, MO, and RD) were obtained by transferring the individual images to the FA template space. See Supplementary Figure 1 for an overview of our image processing procedures.

#### Functional annotation of genetic signals

We obtained cell-type-specific active chromatin annotations per SNP from Finucane et al. (2015) and Boyle et al. (2017). According to Finucane et al. (2015), we performed functional annotation analyses using cell-type-specific annotations marked by four histones: H3K4me1, H3K4me3, H3K9ac and H3K27ac. Each cell-type-specific annotation corresponded to a histone mark in a single cell type, and there were 220 such annotations. The 220 cell-type-specific annotations were further divided into ten groups, including adrenal gland and pancreas, CNS, cardiovascular system, connective tissue and bone, gastrointestinal, immune and hematopoietic systems, kidney, liver, skeletal muscle and other. In our DTI analysis, the SNPs were further divided into three nonoverlapping groups according to their activeness in all cell-type groups and CNS group. A SNP was labeled "CNS_active" if it was annotated as active in CNS cell-type group, and a SNP was labeled "CNS_inactive" if it was annotated as inactive in CNS group, but was active in at least one of the other cell-type groups, and SNPs that were not active in any cell type were labeled "Always_inactive".
